# Genome-wide association analysis links multiple psychiatric liability genes to oscillatory brain activity

**DOI:** 10.1101/232330

**Authors:** Dirk JA Smit, Margaret J Wright, Jacquelyn L Meyers, Nicholas G Martin, Yvonne YW Ho, Stephen M Malone, Jian Zhang, Scott J Burwell, David B Chorlian, Eco JC de Geus, Damiaan Denys, Narelle K Hansell, Jouke-Jan Hottenga, Matt McGue, Catharina EM van Beijsterveldt, Neda Jahanshad, Paul M Thompson, Christopher D Whelan, Sarah E Medland, Bernice Porjesz, William G Iacono, Dorret I Boomsma

**Affiliations:** Psychiatry department, Amsterdam Neuroscience, Academic Medical Center, University of Amsterdam, The Netherlands; Queensland Brain Institute, University of Queensland, Brisbane, Australia; Centre of Advanced Imaging, University Queensland, Brisbane, Australia; Henri Begleiter Neurodynamics Lab., Department of Psychiatry, State University of New York Downstate Medical Center, Brooklyn, New York, USA; QIMR Berghofer Medical Research Institute, Brisbane, Australia; Department of Psychology, University of Minnesota, Minneapolis, MN, USA; Biological Psychology, Amsterdam Public Health research institute, Vrije Universiteit Amsterdam, The Netherlands; Imaging Genetics Center, USC Mark and Mary Stevens Neuroimaging and Informatics Institute, Keck School of Medicine of University of Southern California, Marina del Rey, CA, USA

## Abstract

Oscillations in neuronal activity are widely thought to play a crucial role in information processing and cortical communication ^1–8^. Brain oscillations have been widely investigated as biomarkers of psychiatric disorders and variation in normal human behavior ^9, 10^, including intelligence ^11, 12^, schizophrenia ^13, 14^, attentional deficits ^15, 16^, and substance use ^17, 18^. Beyond the biomarker, oscillatory activity may indeed cause variation in behavior, as it has been shown that disrupting oscillatory activity by blocking GABAergic fast-spiking interneurons in the frontal cortex of mice impairs behavioral flexibility^8^, consistent with observed aberrant frontal cortical oscillatory responses during cognitive tasks requiring cognitive flexibility in humans^19^. Moreover, restoring oscillatory activity in Dlx5/6+/− transgenic mice with similar behavioral impairment—via optogenetic driving of GABA interneurons at gamma frequencies—restored normal behavioral flexibility, thus further supporting the causal involvement of oscillations in driving behavior ^8^.

As one of the most heritable traits in humans ^20–25^, studies have attempted to link oscillatory activity to the genetic liability psychiatric disorders via twin and family studies. However, studies linking specific genetic variants for oscillation strength to psychopathology remain scarce. Here, we performed a meta-analysis of genome wide association studies (GWAS) between genetic variants and the oscillation strength in different frequency bands of electroencephalographic (EEG) recorded activity. In addition, we conducted gene-based analyses, compared these to known liability genes for psychiatric disorders, and examined gene-expression pathways.

The largest study of EEG power and peak frequency to date ^26^ did not yield genome-wide significant hits, but reported a significant contribution of common variants to heritability using random-effects modeling ^27^. In other studies, the most consistent finding has been the involvement of GABA functioning in ~20 Hz beta oscillatory activity. Porjesz et al. ^28^ showed significant linkage between beta oscillations on chromosome 4 and the GABRB1 microsatellite marker, which was overlying a cluster of GABA_A_ receptor genes: GABRG1, GABRA2, GABRA4 and GABRB1. Regional SNP association analysis subsequently pointed to SNPs intronic to GABRA2 as accounting for the signal ^29^. GABRA2 was subsequently associated with both beta oscillations and alcohol use disorders ^29, 30^. A recent study conducted in 117 families of European ancestry from the Collaborative Study on the Genetics of Alcoholism associated several intergenic SNPs at 6q22 with >20 Hz fast beta oscillations ^31^. One of the main aims of the current study is to investigate how genetic variation in GABRA2 affects expression in brain tissues, extend to other oscillations frequencies, and whether liability genes for other psychopathological conditions influence brain function.

Since the effects of single genetic variants on brain activity are expected to be small, GWAS requires large sample sizes. Increases in sample size and statistical power may be obtained by meta-analysis of separate genome-wide association studies (GWAS). The current study describes the results from the ENIGMA-EEG workgroup of the ENIGMA consortium ^32, 33^. We developed EEG processing protocols to extract common measures for band power in the standard frequency bands delta, theta, alpha and beta power at the vertex (Cz) electrode, and occipital (O1, O2) alpha power and alpha peak frequency consistent with ^26^. GWAS results of three population-based and two alcohol-dependence ascertained twin and family cohorts from the Netherlands, Australia, and US were combined in a meta-analysis for a total of 8425 individuals (see also ^34^).

To gain further understanding of the results, extensive follow-up analyses were conducted. We performed enrichment analysis of expression Quantitative Trait Loci (eQTLs) in GTEx brain tissues, positional gene-based analysis, and SNP-based co-heritability analysis of our six EEG traits with related brain phenotypes and psychiatric traits. Follow-up analyses identified genetic variants previously implicated in schizophrenia, including eQTLs that influence gene expression in frontal cortical, anterior cingulate, and subcortical tissues. Hippocampal GABRA2 expression was linked to beta oscillations.

## RESULTS

### Genome-wide association

Supplementary figure S1 shows the Manhattan plots for the six EEG traits. Genome-wide significance was set at 5·10^−8^. Two SNPs were genome-wide significant for Cz alpha power: (rs984924, p=4.7·10^−8^ and rs10231372, p=2.910^−8^); rs984924 on chromosome 4 is an intronic variant within protein kinase cGMP-dependent type II (PRKG2), and rs10231372 on chromosome 7 is an intronic variant within the long non-coding RNA gene LINC00996. Suggestive peaks (p<5·10^−7^) were found for Cz delta power on chromosome 5 (rs6867021, p=1.1·10^−7^), chromosome 6 (rs17055223,p=3.1·10^−7^), chromosome 2 (rs11677128, p=4.3·10^−7^); Cz alpha power on chromosome 1 (rs10910665, p=1.8·10^−7^) and on chromosome 13 (rs9514041, p=1.4·10^−7^). Supplementary Table 1 shows the genome-wide significant SNPs, suggestive peaks, and FDR significant discoveries.

Q-Q plots for the meta-analysis are provided in Figure 1 (pink dots). Full genome median lambdas ranged from 1.02 to 1.06. Subsequent LD score-intercepts—which can be used to parse inflation into true and spurious effects—were not significant (abs(z)<1.50); however, beta power did show significant inflation (z=2.1). Correction of p-values using the intercept had only minor effect and did not change any SNP or gene-based results. Overall, LD score intercept results indicated that there is no evidence of substantial inflation of statistics due to, for example, residual population stratification effects. *Positional gene-based analysis* We performed gene-based analysis using KGG Extended Simes test for each of the EEG traits, which combined the SNP p-values within genes plus flanking regions 50k basepair extensions in 5’ and 3’ UTR directions while taking into account the LD structure. Q-Q plots are included in Figure 1 (blue triangles). Plots showed inflation for gene p-values compared to SNP p-values. Figure 2 shows the gene-based Manhattan plots. FDR-corrected p-values showed significant genes for delta, theta, and alpha power at the vertex. Supplementary Table 2 shows the statistically significant gene discoveries (FDR q=0.05).

**Figure 1.**
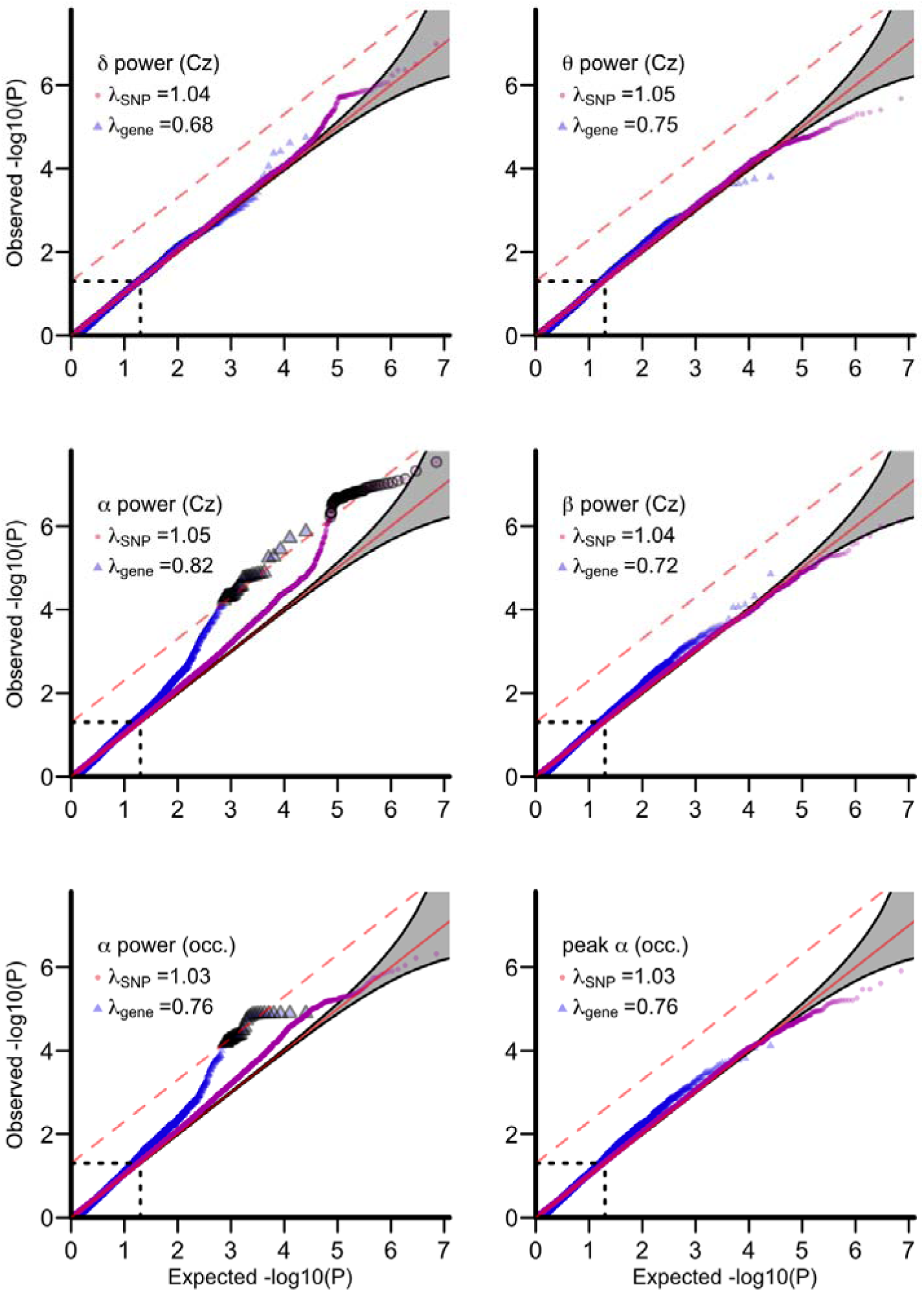
*Quantile-quantile plots of observed versus expected –log_10_(p) for delta, theta, alpha, and beta EEG power at the vertex, occipital alpha power, and occipital alpha peak frequency. Red line is the expected null, grey area is the 95% confidence interval. Dashed red line is the Benjamini-Hochberg FDR q=0.05 threshold. Pink dots are meta-analyzed SNP p-values. FDR-corrected significance is reached for alpha power at Cz. Blue triangles are KGG gene-based test p-values combining SNP effects within gene regions plus 50k base pairs 3’ and 5’ UTR. Many genes reach FDR significance for alpha oscillation power (Supplementary Table 2)*.

**Figure 2.**
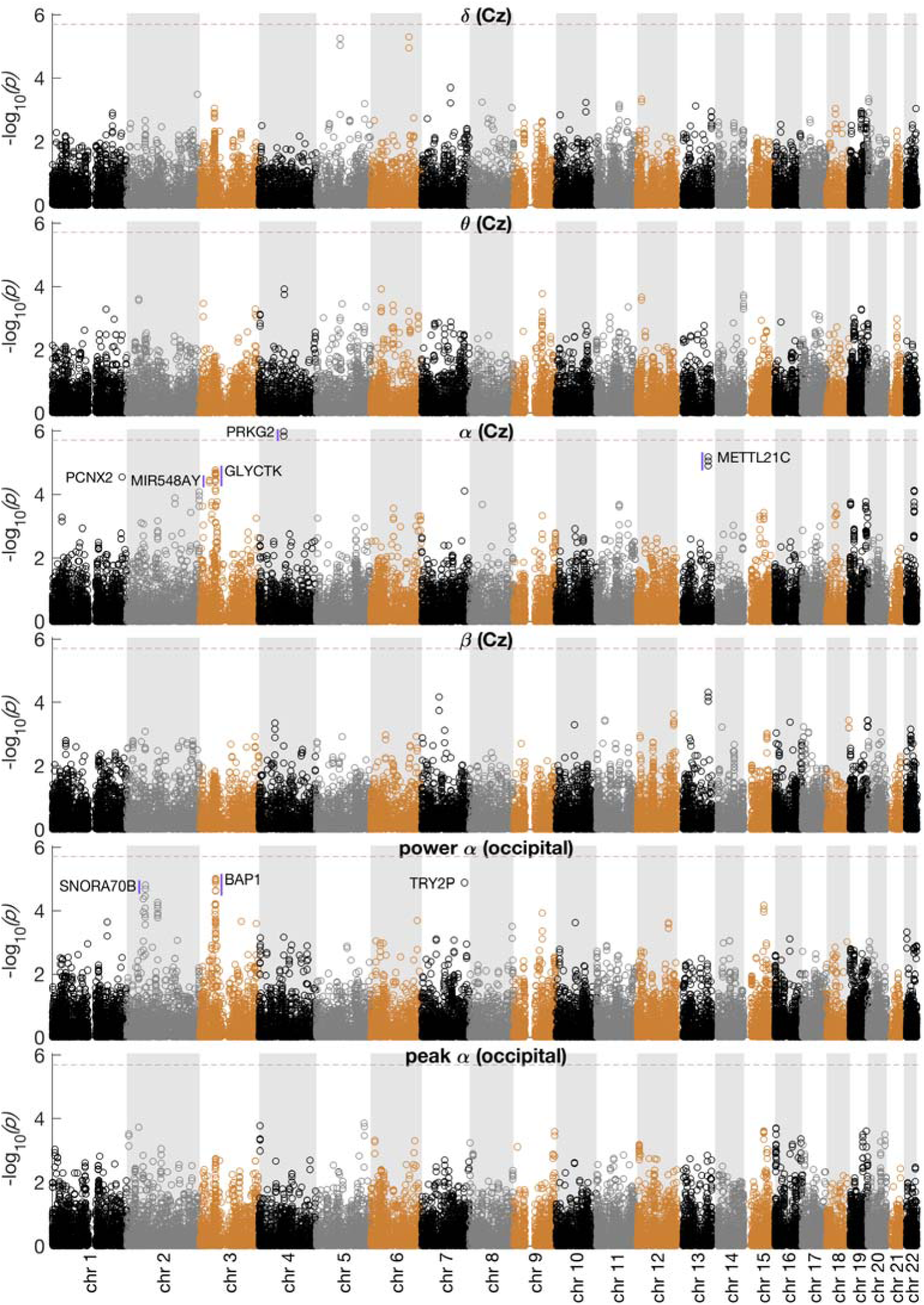
*KGG gene-based test results Manhattan plots for the six EEG traits measured at the vertex electrode (Cz) and occipital (O1/O2). Dashed line is the threshold for genome-wide significance. Named genes are significant discoveries under FDR q=0.05. Only peak findings in the significant region marked with blue vertical lines are shown. A full listing of FDR significant genes is provided in Supplementary Table 2*.

Consistent with the SNP-based findings, PRKG2 was significant for Cz alpha power (p=0.019). LOC101928942 (p=0.019) is an antisense noncoding RNA gene embedded in PRKG2. Both Cz and occipital alpha power showed a cluster of significant genes at 3p21 ranging from (hg19) basepair positions 52234203 to 52728499 (ALAS1 to GLT8D1). For Cz alpha power, 17 of these genes were significant discoveries at q=0.05. For occipital alpha power, 11 genes reached significance of which 4 overlapped with Cz alpha. Figure S2 (top) shows the regional Cz alpha LocusZoom plot of the chromosome 3 region revealing high LD from about 52.2 to 52.8 Mb (hg19). Variants within the same region have been consistently associated with schizophrenia and bipolar disorder ^35, 36^. Figure S2 (bottom) shows the regional association plot for the second Psychiatric Genomics Consortium schizophrenia GWAS ^37^ for comparison. Top SNP in this region was rs7614727 (p=2.0×10^−6^), which is intronic to *WDR82*—previously associated with bipolar disorder and schizophrenia.

Further significant findings were a cluster of three genes (METTL21C, TPP2, and CCDC168; FDR p=0.033 for all) for Cz alpha power; and at 2p15 for occipital alpha power.

### eQTL Expression analysis

To investigate which genes are likely to mediate phenotypic variation in high LD regions and to elucidate mediating brain tissue expression pathways, we performed eQTL analysis. A substantial percentage of eQTLs affect the expression of genes at a distance, often including variants close to a different gene in a different LD region ^38^. *Cis*-eQTLs are genetic variants within a 1Mb region of a gene that explain variability in the expression of the gene in a target tissue ^39–41^. We selected eQTLs from eight brain tissues from the GTEx database ^41^. Cz alpha power associated p-values resulted in inflated Q-Q plots for all tissues (Figure S3). Benjamini-Hochberg FDR significant effects at q=0.05 were observed for the Frontal Cortex and Anterior Cingulate Gyrus, and the Hypothalamus. Occipital alpha power showed similar effects, but for different brain regions (Caudate, Nucleus Accumbens, Hippocampus) (Figure S4). The significant SNPs were frontal cortical tissue eQTLs for MTERF4, GNL3, and ITIH4; the latter two being schizophrenia/bipolar disorder liability genes at 3p21. Significant SNPs for occipital power were cortical-tissue eQTLs for genes IL1RL1, IL18R1, CLHC1, GLYCTK, and ITIH4.

To test for overall significance of gene-expression enrichment in alpha oscillation power, we used the online tool FUMA ^42^. We extracted the top 500 genes from the gene-based association (Cz and occipital alpha), which were matched against genes significantly up- or downregulated in each GTEx tissue compared to the average of other tissues (i.e., differentially expressed genes determined by a Bonferroni-corrected t-test). Significance of enrichment was determined by the hypergeometric test with Bonferroni correction. Figure 3 shows that brain derived tissues are almost invariably significant, and much more so than other tissues.

**Figure 3.**
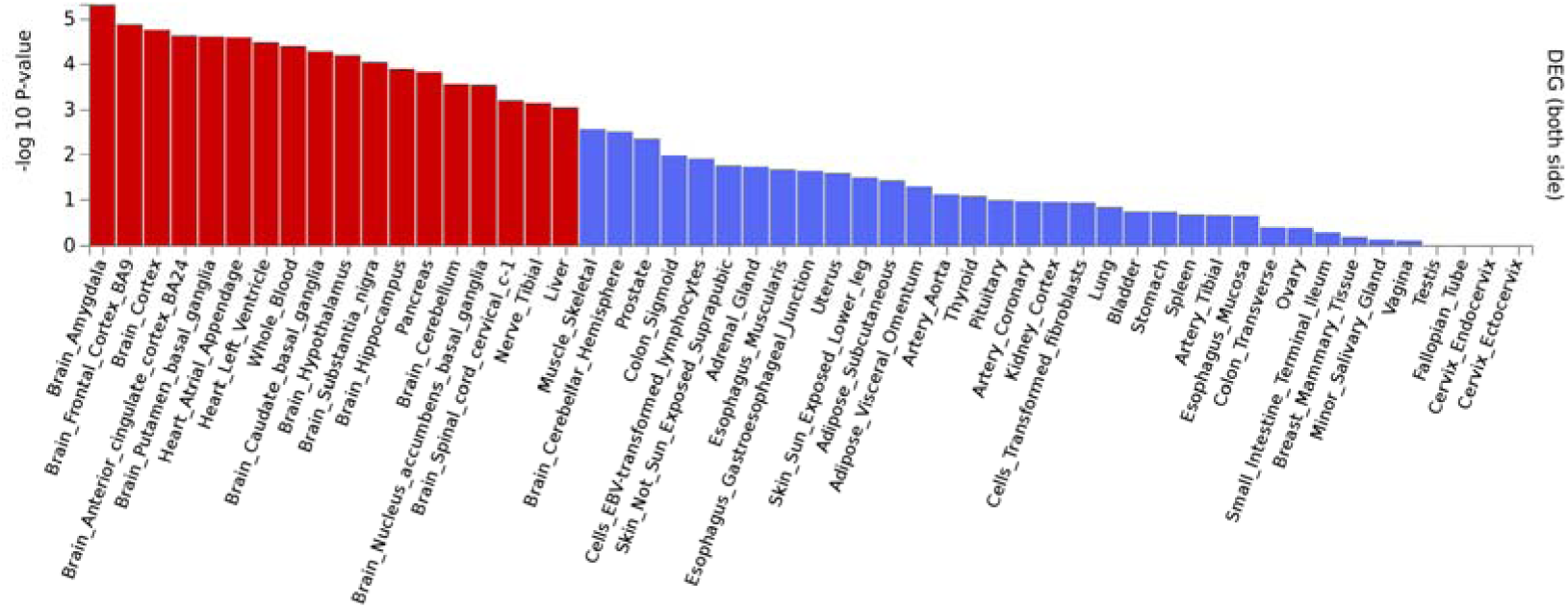
*FUMA^42^ enrichment analysis. The top 500 genes in the occipital alpha analysis are significantly enriched for genes that are up or down regulated in all cortical and subcortical brain tissues from the GTEx database, save cerebellar hemisphere. Other significantly enriched tissues are heart, whole blood, pancreas, tibial nerve, and liver*.

### Imputed gene-expression association of alpha oscillations

To further elucidate tissue-expression pathways of the chromosome 2 cytokine receptor genes and the schizophrenia liability genes on chromosome 3 affecting alpha oscillation power we applied MetaXcan using the all GTEx brain tissues ^40, 43^. MetaXcan may have increased power to detect significant gene / phenotype associations by combining genetic variants in a sparse elastic net prediction model. Four genes near the 3p21 region showed at least one FDR significant association (ITIH4, GNL3, GLYCTK, and TEX264). Figure 4 shows the expression profiles for chromosome 3 genes with at least one FDR significant effect. ITIH4 showed a more widespread association across tissues (significant in hypothalamus), whereas GLYTCK showed a rather specific hippocampal expression for both occipital and Cz alpha power. Immune genes IL1RL1 and IL18R1 on chromosome 2 also reached the threshold for significance in the significant association with cortical and subcortical expression with alpha oscillations (see Figure 4).

**Figure 4.**
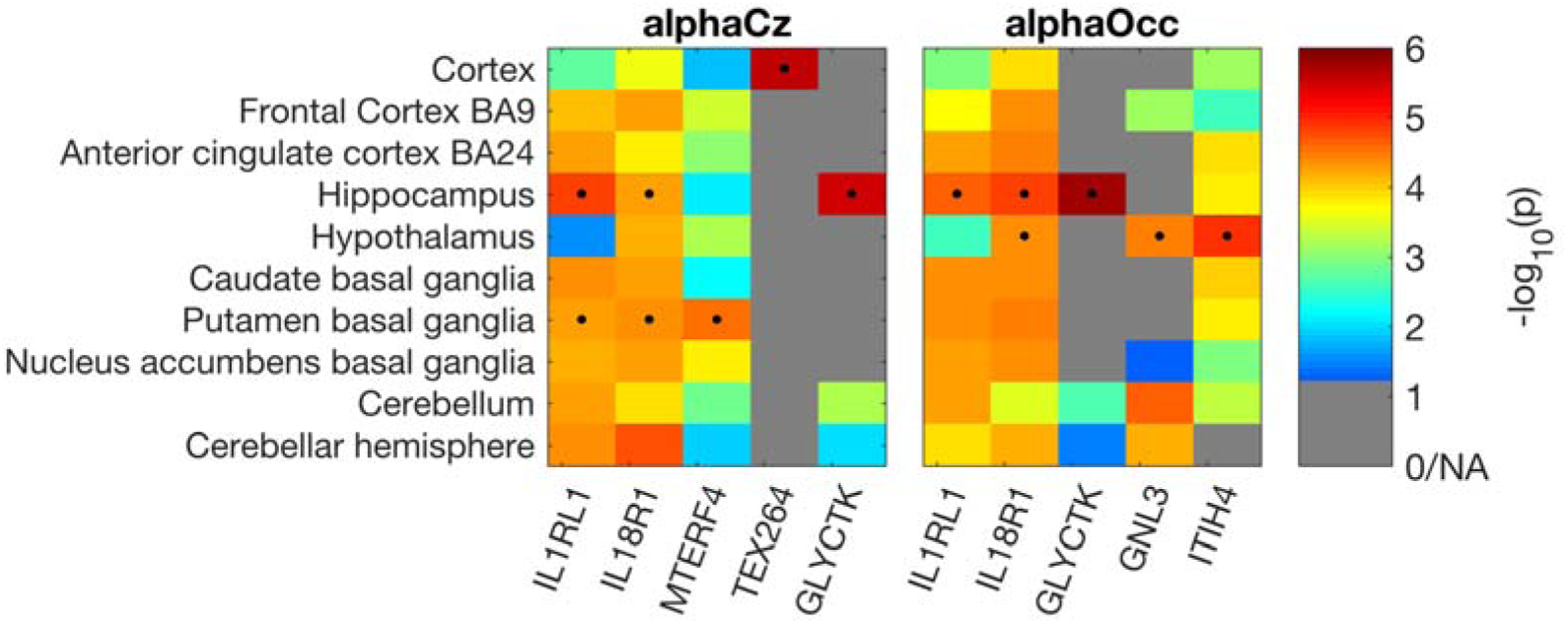
MetaXcan gene-expression results for chromosomes 2 and 3 indicate which positional gene-based discoveries and which brain tissues may be involved in affecting alpha oscillation power. Uncorrected p-values are shown, FDR significant discoveries are marked with a dot. Out of the 24 significant genes in region 3p21, GLYCTK associated with alpha power via hippocampal expression, TEX264 via general cortical expression, GNL3 and ITIH4 via hypothalamic expression. Furthermore, IL18R1, IL1RL1, and MTERF4 on chromosome 2 were not discovered in the positional analysis. These genes affected alpha oscillations via the Putamen and Hippocampus, but IL1RL1 and IL18R1 expression in many other tissues just failed to reach the FDR threshold. Note that FDR correction was performed within tissue.

### GABA_A_ receptor genes

Planned comparisons for two GABA_A_ receptor genes at 4p12 were made in relation to beta power. The results from the KGG GATES positional gene-based analysis showed p=0.024 for GABRA2 and p=0.15 for GABRB1. After removing the results from the COGA sample—on which the previous positive findings were based—the p-value increased to p=0.052 for GABRA2. MetaXcan analysis showed that imputed expression was associated with beta power for hippocampal expression of GABRA2 (p=0.0024) and not with other tissues (p>0.2).

### SNP-based heritability

LD score regression for Hapmap 3 annotated SNPs (N_SNP_>819k) was carried out for SNP effects from the meta-analysis. Heritability estimates (Figure S5) were lower than estimates from twin and family studies of EEG power (ranging from 0.45 to 0.9; e.g., see ^21^), but are in line with the GCTA random-effects modeling on common SNPs for the same EEG measures ^26^.

### Genetic correlation analysis

Bivariate LD score regression was used to calculate genetic correlation r_G_ between traits. Figure S6 shows the genetic correlations between the EEG phenotypes. Strong and positive genetic correlations were observed among the EEG power phenotypes. Occipital and Cz alpha power correlated 1.0, and correlated highly with beta power (r_G_=.67 and r_G_=0.71 respectively). Slow oscillatory power (delta and theta) also correlated near 1.0. Only the theta with beta power genetic correlation was modest and not statistically significant. Negative correlations between peak alpha frequency and the slow oscillation power phenotypes were observed, and a nonsignificant positive correlation with beta power.

LD score regression based genetic correlations with a wide range of human phenotypes with published GWAS results are shown in Figure S7. Significant effects were observed but failed to survive multiple testing correction (FDR or Bonferroni), which may be expected in light of the still relatively low sample size of the current GWAS and the subsequently large confidence intervals. Top effects included r_G_=−0.35 (uncorrected p=0.0094) between theta power and autism spectrum disorder, and r_G_=0.55 (uncorrected p=0.014) between beta power and generalized epilepsy.

## DISCUSSION

We have presented results from the first international consortium for investigating the molecular genetic basis of brain functional activity as measured by resting EEG. The results revealed two genome-wide significant associations for Cz alpha power: on chromosome 4, a SNP intronic to PRKG2 (rs984924), and on chromosome 7, a SNP intronic to LINC00996 (rs10231372). FDR correction yielded 68 significant SNPs in the same PRKG2 and LINC00996 regions, plus intronic variants within PCNX2 and METTL21C. Gene-based analyses identified multiple genes significantly associated with Cz and occipital alpha power, including PRKG2, METTL21C, and several genes in a region on chromosome 3. PRKG2 influences anthropometric and blood pressure-related traits ^44,45^ and also affects multiple phenotypes in mice, including skeletal and adipose tissues. Humans with 4q21 microdeletion syndrome—which includes PRKG2 and flanking genes—show similar skeletal symptoms, including facial bone and growth retardations, but also neuropsychological symptoms, including speech and mental retardation ^46, 47^.

KGG gene-based analyses implicated a high-LD region on chromosome 3 that included many significant genes associated with Cz and occipital alpha power. Variants in this region have been associated with schizophrenia and bipolar disorder. Significant brain-tissue eQTLs pointed to ITIH4, GNL3, and GLYCTK as genes with altered expression. MetaXcan significantly associated widespread brain expression for ITIH4, with hypothalamic expression reaching significance. For GNL3, hypothalamic and cerebellar tissues significantly associated with alpha oscillations. Hippocampal GLYCTK expression associated with Cz and occipital alpha. MetaXcan further associated cortical TEX264 expression. By using expression analyses, we were able to strongly reduce the number of target genes in the chromosome 3 region from twenty-four to four, and localize their effects to hypothalamic and hippocampal expression as most strongly associated with alpha oscillations.

The association schizophrenia liability genes with oscillatory brain activity and the specific tissues with significantly altered expression highlights where oscillatory brain activity changes with increased disease risk. Altered expression of ITIH4 in the frontal cortex in the context of schizophrenia has recently been reported ^48^, and is consistent with reduced alpha oscillatory activity in the frontal cortex observed in schizophrenia ^13,49–52^. FDR significant SNPs were eQTLs for ITIH4, GLN3, and MTERF4 in the frontal cortex for Cz alpha oscillations. Our results indicate that schizophrenia liability gene ITIH4 affects oscillatory brain function, and adds GNL3 and MTERF4 as possible target genes. Brain eQTLs further pointed to cytokine receptor genes IL1RL1 and IL18R1, which are immune system genes linked to asthma, celiac disease, IBS, and atopic dermatitis ^53–56^. MetaXcan imputed expression analysis indicated that these genes are also brain expressed, and associated with alpha oscillation power for widespread cortical and subcortical tissues, reaching significance for the Hippocampus and Putamen. The association of IL18R1 expression with schizophrenia was reported recently ^57^. Our results indicate that these immunological liability genes also affect oscillatory brain function by altering widespread expression in the brain.

Results confirmed GABA_A_ signaling as being involved in fast oscillatory (beta) activity in the full meta-analysis (p=0.024). Expression analysis significantly associated hippocampal GABRA2 expression to beta oscillations (p=0.0024). This result fits well with observations that beta oscillations are influenced by GABAA receptor \alpha 2 agonists such as benzodiapines ^58–60^, and the crucial role of GABA_A_ interneurons for synchronized fast rhythms in the brain ^61^. In our view, there is now strong evidence that GABA mediates the relation between resting-state EEG beta power and alcohol dependence ^17, 29, 62^. The selective hippocampal expression association suggests that the genetic variants affecting beta oscillations also affect hippocampal GABA_A_ receptor’s sensitivity to interneuron inhibition.

Twin and family studies have consistently indicated that EEG alpha power is one of the most heritable traits in humans at up to 96% for frontal alpha power in young adult samples ^20, 21, 24, 63^. The SNP heritability observed here using LD score regression was only able to retrieve a relatively small proportion of variance of the often highly heritable EEG traits. This discrepancy could be caused by a relative large contribution of rare SNPs that are poorly tagged by the common SNP arrays used here. The SNP-based genetic correlation analysis was more consistent with twin/family studies. Strong genetic correlations (>.70) were observed among the slower (delta theta) and among faster oscillations (alpha beta). Results from twin studies generally ranged from 0.50 to 0.90, although the twin-based r_G_ between theta and delta oscillation power is generally not as strong as the SNP r_G_ observed here. This inconsistency between twin/family and SNP coheritability could perhaps be explained by the restricted scalp locations tested in the current analysis and/or the sample heterogeneity (e.g., COGA selected for alcohol use disorders).

Coheritability analysis showed a nominally significant genetic correlation (r_G_=0.55) between beta oscillations and generalized epilepsy. This is consistent with the putative role of fast beta/gamma oscillations in ictogenesis (in the present sample only nonaffected individuals were used). The largest epilepsy GWAS to date found suggestive evidence for the involvement of GABRA2 ^64^, which we found to be related to beta oscillation power. GABA is a main antiepileptic drug target, and is known to affect (motor) beta EEG via GABAergic modulation of pyramidal cells ^65–68^. We additionally observed a significant genetic correlation between autism and theta oscillations (r_G_=−0.35, p=0.009). Although deviant brain function in autism is most consistently found in the lower gamma band due to altered GABA inhibitory neuronal action ^69–72^, cortical and hippocampal theta/lower alpha are known to show phase-amplitude coupling with fast oscillations (gamma ^73–75^). The significant genetic correlation (r_G_=0.22, p=0.019) between heart rate and alpha power is consistent with observations in concurrent EEG and ECG recordings ^76^.

In sum, we have found evidence that (hippocampal) GABA receptor alpha 2 subunit is involved in altering beta power, possibly via hippocampal expression—consistent with its relation to epilepsy and alcohol dependence that are both well known for the involvement of GABA_A_ ^29, 77^. Schizophrenia liability genes on chromosome 2 and 3 affected alpha oscillation power. SNPs in the chromosome 3 region were eQTLs for ITIH4, GNL3, and GLYCTK. Significant eQTLs were tissue specific, including the frontal cortex, anterior cingulate cortex, hypothalamus, and hippocampus. Expression analysis further targeted immune system genes IL1RL1 and IL18R1 with altered expression in Putamen and Hippocampus.

GWAS is dependent on very large sample sizes as the effects of the tagging genetic variants (SNPs) are small, even for brain endophenotypes ^78, 79^. Our results suggest that EEG power GWAS may indeed be somewhat more powerful than of complex behavioral phenotypes, which generally require larger sample sizes. Current sample sizes may be adequate to reveal significant individual genetic associations, pathways for the expression of psychiatric liability in the brain, and prove hopeful for future GWAS of additional EEG parameters. For example, two recently published GWASs of bipolar EEG from families of African and European ancestry reported genome-wide signal at 3q26 and 6q22, respectively ^31, 80^. Bipolar EEG derivations show more localized activity than other EEG derivations and remove volume conduction effects, and have been particularly successful as a biomarker of alcohol dependence. Other EEG parameters of high interest are functional connectivity as biomarker for various neurodevelopmental and psychiatric disorders. The current results indicate that finding genetic variants or genes related to these EEG parameters is entirely feasible.

## METHODS

### Subjects

Resting state EEG and genome-wide genotyping were available for a total of 8,425 individuals from Australia, the Netherlands, and the USA. All subjects were part of twin and family studies into the genetics of health and behavioral traits with additional psychophysiological assessments: the Minnesota Twin Family Study (MTFS), the Collaborative Study on the Genetics of Alcoholism (COGA), the Brisbane Adolescent Twin Study (BATS), and the Netherlands Twin Register (NTR). All sites excluded subjects with a history of neurological problems, including tumor and head trauma. Alcohol dependence ascertained samples (COGA EA and COGA case control) required alcohol testing before EEG recording. Full details and demographics for each cohort are given in the supplementary information.

### EEG recording and preprocessing

For details on EEG assessment by the individual groups see supplementary material.

### EEG Power and Peak frequency analysis

All groups analyzed the vertex recordings (Cz) and the average of Occipital leads (O1, O2) in standard frequency bands. Cleaned data were imported into MATLAB, epoched into 2s epochs, and power spectra calculated using FFT. Frequency bins were defined as delta (1-3.75Hz), theta (4-7.75 Hz), alpha (8-12.75Hz), and beta (15-25 Hz). Power was defined as the squared radius of the orthogonal sine and cosine amplitudes averaged over window size, and the mean value taken for the frequency band to obtain power density. Alpha peak frequency was determined using the power-weighted method in accordance with ^26^ between 7 and 14 Hz.

### QC and Genome-wide association analysis

Groups used dosage based imputed SNP sets using CEU reference panels hg19/build 37 from 1000 genomes phase1 or phase3. Imputation followed ENIGMA imputation protocols ^32^ (http://enigma.usc.edu/wp-content/uploads/2012/07/ENIGMA21KGPcookbookv3.pdf). Association analyses accounted for family relatedness. Sex, Age and age^2 were used as covariates plus Principal Components plus disease status (when applicable). See supplementary materials for more group specific methods and/or deviance from these standard analyses protocols, and covariate analysis results.

Pre-meta-analysis QC were performed using EasyQC ^81^. We filtered on sample MAF (0.03 for the largest dataset, MTFS; 0.04 for the intermediate datasets, COGA case control and QIMR BATS; and 0.05 for NTR and COGA EA), as well as EUR 1000 Genomes reference set MAF (<0.03), N<200, HWE p<10^−7^, INFO<0.8, INFO>1.05, imputation R^2^<0.4, invalid numbers (Inf, NA), 0.2 difference between sample and reference set allele frequencies. Further checks consisted of matching alleles, duplicates, and strand flips. Meta-analyzed SNPs were filtered for combined N>6000. The final datasets consisted of 4959085 to 4959521 SNPs depending on phenotype.

### SNP heritability

LD score regression ^82^ uses the natural experiment present in the genome due to variable amounts of Linkage Disequilibrium (LD) between SNPs. Causal variants will cause straight slope decline in test statistics of nearby SNPs with decreasing levels of LD to the causal variant in the case of additive genetic variation. The slope of the regression line of the chi-square statistic against LD scores across the genome reflects the heritability of the trait ^82^. SNP heritability estimation using LD score regression has the advantage of being insensitive to population stratification effects, as these will result in an upward shift across all LD score bins, thus affecting the intercept and not the slope of the LD-dependent regression.

We used LD score regression to estimate the SNP based heritability of the six phenotypes following the recommendations in ^82^, including pruning for Hapmap 3 SNPs. Next, the LD-score regression intercept was used to assess quality of the GWAS and removal of stratification effects by the population Principal Components (see for example ^83^). Finally, we used bivariate LD-score regression to estimate genetic correlation r_G_, between the EEG phenotypes and GWASs available in LD Hub (http://ldsc.broadinstitute.org) ^84, 85^. This includes GWASs on schizophrenia, bipolar disorder, subjective wellbeing, and neuroticism, and ENIGMA subcortical volumes, intracranial volume, and brain volume (See Supplementary Methods for a full reference list). We extended these with generalized epilepsy and educational attainment ^64, 86^.

## ACKNOWLEDGEMENTS

This research was carried out under the auspices of the Enhancing Imaging Genetics through Meta-Analysis (ENIGMA) consortium. The ENIGMA-EEG Working Group gratefully acknowledges support from the NIH Big Data to Knowledge (BD2K) award (U54 EB020403 to Paul Thompson). Summary statistics of the GWASs will be made available via the ENIGMA website (http://enigma.ini.usc.edu).

<u>The Collaborative Study of the Genetics of Alcoholism (COGA)</u> continues to be inspired by our memories of Henri Begleiter and Theodore Reich, founding PI and Co-PI of COGA, and acknowledges all participating centers (see supplementary note). We gratefully acknowledge the following funding sources: <u>University of Minnesota:</u> Funding by National Institutes of Health (NIH) DA 05147, DA 36216, DA 024417. <u>COGA</u>: NIH NIAAA U10AA00840, NIH GEI U01HG004438, HHSN268200782096C. <u>Vrije Universiteit</u>: NOW/ZonMW 904-61-090, 985-10-002, 912-10-020, 904-61-193,480-04-004, 463-06-001, 451-04-034, 400-05-717, Addiction-31160008, Middelgroot-911-09-032, Spinozapremie 56-464-14192, BBMRI–NL, 184.021.007, NWO-Groot 480-15-001/674. ERC FP7/2007-2013), ENGAGE HEALTH-F4-2007-201413, ERC Advanced 230374, Starting 284167), NIMH U24 MH068457-06, NIH R01D0042157-01A, MH081802, R01 DK092127-04, Grand Opportunity grants 1RC2 MH089951 and 1RC2 MH089995. <u>Personal funding</u>: NWO 480-04- 004, NWO/SPI 56-464-14192, and NWO 911-09-032 to D.B., NWO/MagW VENI-451-08-026 to D.S.; ERC-230374 to D.B.; BBR Foundation (NARSAD) 21668 to D.S.; VU-USF 96/22 to D.B.; HFSP RG0154/1998-B to D.B. and E.d.G.; KNAW PAH/6635 to D.B.; Australian Research Council A79600334, A79906588, A79801419, DP0212016 to N.M. and M.W.; NHMRC 389891 to N.M.; Fellowship APP1103623 to S.M.; NIH K01DA037914 to J.L.M. The Genotype-Tissue Expression (GTEx) Project was supported by the Common Fund of the Office of the Director of the National Institutes of Health, and by NCI, NHGRI, NHLBI, NIDA, NIMH, and NINDS. The data used for the analyses described in this manuscript were obtained from the GTEx Portal on 20 Oct 2016. See supplementary note for a full acknowledgement.

## SUPPLEMENTARY NOTE: Full acknowledgement

<u>University of Minnesota:</u> Funding by National Institutes of Health (NIH) DA 05147, DA 36216, DA 024417, AA 09367, DA 13240

<u>COGA:</u> The Collaborative Study on the Genetics of Alcoholism (COGA), Principal Investigators B. Porjesz, V. Hesselbrock, H. Edenberg, L. Bierut, includes eleven different centers: University of Connecticut (V. Hesselbrock); Indiana University (H.J. Edenberg, J. Nurnberger Jr., T. Foroud); University of Iowa (S. Kuperman, J. Kramer); SUNY Downstate (B. Porjesz); Washington University in St. Louis (L. Bierut, J. Rice, K. Bucholz, A. Agrawal); University of California at San Diego (M. Schuckit); Rutgers University (J. Tischfield, A. Brooks); Department of Biomedical and Health Informatics, The Children’s Hospital of Philadelphia; Department of Genetics, Perelman School of Medicine, University of Pennsylvania, Philadelphia PA (L. Almasy), Virginia Commonwealth University (D. Dick), Icahn School of Medicine at Mount Sinai (A. Goate), and Howard University (R. Taylor). Other COGA collaborators include: L. Bauer (University of Connecticut); J. McClintick, L. Wetherill, X. Xuei, Y. Liu, D. Lai, S. O’Connor, M. Plawecki, S. Lourens (Indiana University); G. Chan (University of Iowa; University of Connecticut); J. Meyers, D. Chorlian, C. Kamarajan, A. Pandey, J. Zhang (SUNY Downstate); J.-C. Wang, M. Kapoor, S. Bertelsen (Icahn School of Medicine at Mount Sinai); A. Anokhin, V. McCutcheon, S. Saccone (Washington University); J. Salvatore, F. Aliev, B. Cho (Virginia Commonwealth University); and Mark Kos (University of Texas Rio Grande Valley). A. Parsian and M. Reilly are the NIAAA Staff Collaborators.

We continue to be inspired by our memories of Henri Begleiter and Theodore Reich, founding PI and Co-PI of COGA, and also owe a debt of gratitude to other past organizers of COGA, including Ting-Kai Li, P. Michael Conneally, Raymond Crowe, and Wendy Reich, for their critical contributions. This national collaborative study is supported by NIH Grant U10AA008401 from the National Institute on Alcohol Abuse and Alcoholism (NIAAA) and the National Institute on Drug Abuse (NIDA). Funding for GWAS genotyping, which was performed at the Johns Hopkins University Center for Inherited Disease Research, was provided by NIAAA, the NIH GEI (U01HG004438) and the NIH contract ‘High through-put genotyping for studying the genetic contributions to human disease’ (HHSN268200782096C).

<u>Netherlands Twin Registry/Vrije Universiteit</u>: Funding by Netherlands Organization for Scientific Research (NWO) and The Netherlands Organisation for Health Research and Development (ZonMW) grants 904-61-090, 985-10-002, 912-10-020, 904-61-193,480-04-004, 463-06-001, 451-04-034, 400-05-717, Addiction-31160008, Middelgroot-911-09-032, Spinozapremie 56-464-14192, Biobanking and Biomolecular Resources Research Infrastructure (BBMRI -NL, 184.021.007, NWO-Groot 480-15-001/674). VU Institute for Health and Care Research (EMGO+); the European Community’s Seventh Framework Program (FP7/2007-2013), ENGAGE (HEALTH-F4-2007-201413); the European Research Council (ERC Advanced, 230374, ERC Starting grant 284167), Rutgers University Cell and DNA Repository (NIMH U24 MH068457-06), the Avera Institute, Sioux Falls, South Dakota (USA) and the National Institutes of Health (NIH, R01D0042157-01A, MH081802; R01 DK092127-04, Grand Opportunity grants 1RC2 MH089951 and 1RC2 MH089995). Part of the genotyping and analyses were funded by the Genetic Association Information Network (GAIN) of the Foundation for the National Institutes of Health. Computing was supported by BiG Grid, the Dutch e-Science Grid, which is financially supported by NWO.

<u>Personal funding:</u> Netherlands Organisation for Scientific Research NWO 480-04-004, NWO/SPI 56-464-14192, and NWO 911-09-032 to D.B., NWO/MagW VENI-451-08-026 to D.S.; European Research Council ERC-230374 to D.B.; BBR Foundation (NARSAD) Young Investigator grant 21668 to D.S.; VU University VU-USF 96/22 to D.B.; Human Frontiers of Science Program RG0154/1998-B to D.B., and E.d.G.; KNAW Academy Professor Award (PAH/6635) to D.B.; Australian Research Council (A79600334, A79906588, A79801419, DP0212016) to N.M. and M.W.; National Health and Medical Research Council (Medical Bioinformatics Genomics Proteomics Program 389891) to N.M.; Fellowship APP1103623 to S.M.; NIH K01DA037914 to J.L.M.

